# ePat: extended PROVEAN annotation tool

**DOI:** 10.1101/2021.12.21.468911

**Authors:** Takumi Ito, Kazutoshi Yoshitake, Takeshi Iwata

## Abstract

The ‘ePat’ (extended PROVEAN annotation tool) is a software tool that extends the functionality of PROVEAN: a software tool for predicting whether amino acid substitutions and indels will affect the biological function of proteins. The ‘ePat’ extends the conventional PROVEAN to enable the following two things, which the conventional PROVEAN could not calculate the pathogenicity of these variants. First is to calculate and score the pathogenicity of indel mutations with frameshift and variants near splice junctions. Second is to use batch processing to calculate the pathogenicity of multiple variants into a variants list (VCF file) in a single step. ePat can help extract variants that affect biological functions by utilizing not only point mutations, and indel mutations that does not cause frameshift, but also frameshift, stop gain, and splice variants. These extended features will increase detection rate and improve diagnostic of inherited diseases or associate specific variant to phenotype.

## 1. Introduction

In recent years, improvements in sequencing technology have generated a large amount of information on mutations in patients with inherited diseases or organisms with specific phenotype compared to normal. Mutations such as amino acid substitutions, insertions, deletions, frameshifts, and acquisition of stop codons significantly affect protein function [1] [2] [3] [4] [5] [6].

Computational tools such as Polyphen [7], SIFT [8], PROVEAN [9], and other prediction tools [10] have been developed to search for variants that cause diseases or specific phenotypes, and to quantify the impact of variants on protein function (pathogenicity). PROVEAN made it enable to predict the pathogenicity of variants not limited to single amino acid substitutions but also in-frame insertions, deletions, and multiple amino acid substitutions [11]. However, these tools have two problems: first, they are unable to calculate variants that are considered to have a significant impact on protein function, such as frameshift and stop gain, and second, they cannot calculate the pathogenicity of variants that spread across multiple genes in a single step.

We have developed ‘ePat’ (extended PROVEAN annotation tool), an extension of PROVEAN that solves the above two problems. Unlike existing tools, ePat is able to calculate the pathogenicity of variants near the splice junction, frameshift, stop gain, and start lost. In addition, batch processing is used to calculate the pathogenicity of all variants in a VCF file in a single step.

## 2. Method

With the given reference, a database for SnpEff is created and input vcf file is annotated with SnpEff. We then extract variants that have a HIGH or MODERATE pathogenicity level as a result of the SnpEff annotation. For each row of the VCF file, information of the variants annotated with SnpEff (INFO column) was extracted. With this information, the variants are classified into 5 categories. The first is a variant near the splice junction (splice variants), the second is frameshift, the third is stop gain, the fourth is start lost, and the last is in-frame variants (point mutation or indel mutations that do not cause frameshift).

Variants from category 1 to 4 are given pathogenicity as defined by ePat, and category 5 is given pathogenicity by PROVEAN. The pathogenicity defined by ePat is calculated with the following method. For each amino acid, calculate the pathogenicity when it is replaced by each of the 20 amino acids. The average of these pathogenicity is used as the pathogenicity for that position. The maximum pathogenicity for each position is the pathogenicity of this variants. The details of the algorithm are described below (Figure1).

**Figure 1.**
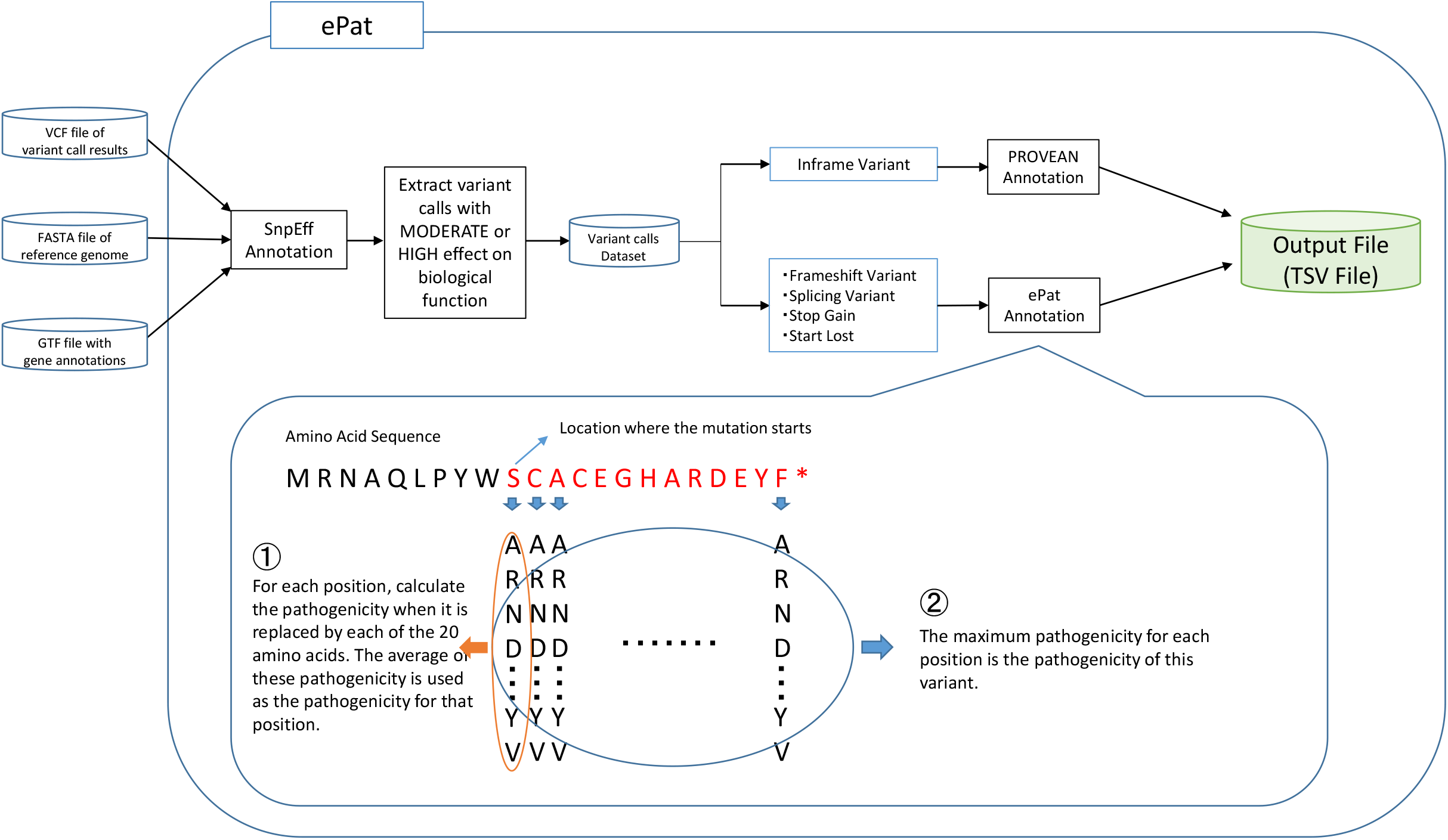
Overview of ePat With the given reference, we create a database for SnpEff and annotate with SnpEff. We then extract variants that have a HIGH or MODERATE pathogenicity level as a result of the SnpEff annotation. For each row of the VCF file, information of the variants annotated with SnpEff was extracted. With this information, the variants are classified into 5 categories. The first is a variant near the splice junction (splice variants), the second is frameshift, the third is stop gain, the fourth is start lost, and the last is in-frame variants (point mutation or indel mutations that do not cause frameshift). Variants from category 1 to 4 are given pathogenicity as defined by ePat, and category 5 is given pathogenicity by PROVEAN. The pathogenicity defined by ePat is calculated with the following method. For each amino acid, calculate the pathogenicity when it is replaced by each of the 20 amino acids. The average of these pathogenicity is used as the pathogenicity for that position. The maximum pathogenicity for each position is the pathogenicity of this variants. The details of the algorithm are described below.

### 2-1. Input file

The input data is a VCF file after variant call (Table 1), a FASTA file of the reference genome, and a GTF file with gene annotations. These input files are given as arguments to the ePat runtime.

**Table 1.** Input overview of ePat

| #CHRO | POS | ID | RE | AL | QUA | FILTE | INFO | FORM | HG0009 |
| --- | --- | --- | --- | --- | --- | --- | --- | --- | --- |
| M |  |  | F | T | L | R |  | AT | 6 |
| 22 | 2469829<br>4 | rs2021421<br>65 | C | G | 100.<br>0 | PASS | AC=1;AF=0.000199681;AN=5008 | GT | 0\|0 |
| 22 | 2470931<br>7 | rs5480176<br>12 | G | A | 100.<br>0 | PASS | AC=1;AF=0.000199681;AN=5008 | GT | 0\|0 |
| 22 | 2470932<br>1 | rs5695942<br>48 | G | C | 100.<br>0 | PASS | AC=1;AF=0.000199681;AN=5008 | GT | 0\|0 |
| 22 | 2470942<br>0 | rs3578391<br>4 | C | T | 100.<br>0 | PASS | AC=4;AF=0.000798722;AN=5008 | GT | 0\|0 |
| 22 | 2470942<br>3 | rs3717804<br>53 | A | G | 100.<br>0 | PASS | AC=1;AF=0.000199681;AN=5008 | GT | 0\|0 |

### 2-2. SnpEff Annotation

With given reference, we create a database for SnpEff and annotate the VCF file with SnpEff. We then extract variants that have a HIGH or MODERATE pathogenicity level as a result of the SnpEff annotation.

### 2-3. Extract Variant Information

For each row of the VCF file, extract the information of the variants annotated with SnpEff ([gene ID, variant type, pathogenicity level, DNA mutation information, amino acid mutation information]) from the INFO column. With this information, the variants are classified into (1) variants near the splice junction (splice variants), (2) frameshift, (3) Stop Gain, (4) Start Lost, and (5) in-frame variants (point Mutation or indel mutations that do not cause frameshift).

### 2-4. Calculate pathogenicity

Variants from (1) to (4) are given pathogenicity as defined by ePat, and those (5) will be given pathogenicity by PROVEAN. The pathogenicity defined by ePat is calculated with the following method.

For each position, calculate the pathogenicity when it is replaced by each of the 20 amino acids. The average of these pathogenicity is used as the pathogenicity for that position. The maximum pathogenicity for each position is the pathogenicity of this mutation.

If the mutation is (1) mutation near splice junctions, then we calculate the pathogenicity defined by ePat in the range from the splice junction where the mutation occurs to the stop codon.

If the mutation is (2) frameshift, then we calculate the pathogenicity defined by ePat in the range from the amino acid where the frameshift starts to the stop codon.

If the mutation is (3) stop codon, then we calculate the pathogenicity defined by ePat in the range from the amino acid to be replaced by the stop codon to the original stop codon. For Stop Lost, the pathogenicity is not calculated.

If the mutation is (4) start lost, then we calculate the pathogenicity defined by ePat in the range from the original start codon to the next methionine.

If the mutation is (5) in-frame variant, then we calculate the pathogenicity by PROVEAN.

## 3. Result and Discussion

After the calculation is completed, the PROVEAN_score and PROVEAN_pred columns are added to the Input file. In the PROVEAN_score column, the pathogenicity calculated by ePat or PROVEAN is described. In the PROVEAN_pred column, if the pathogenicity is less than - 2.5, then D (Damaged) is described, meaning that the mutation is likely to affect amino acid function. If the pathogenicity is greater than -2.5, then N (Damaged) is described, meaning that the mutation is unlikely to affect amino acid function. The output is named as output_provean_{PREFIX_OF_YOUR_INPUTFILE}.txt and saved in the output directory (Table 2).

**Table 2.**
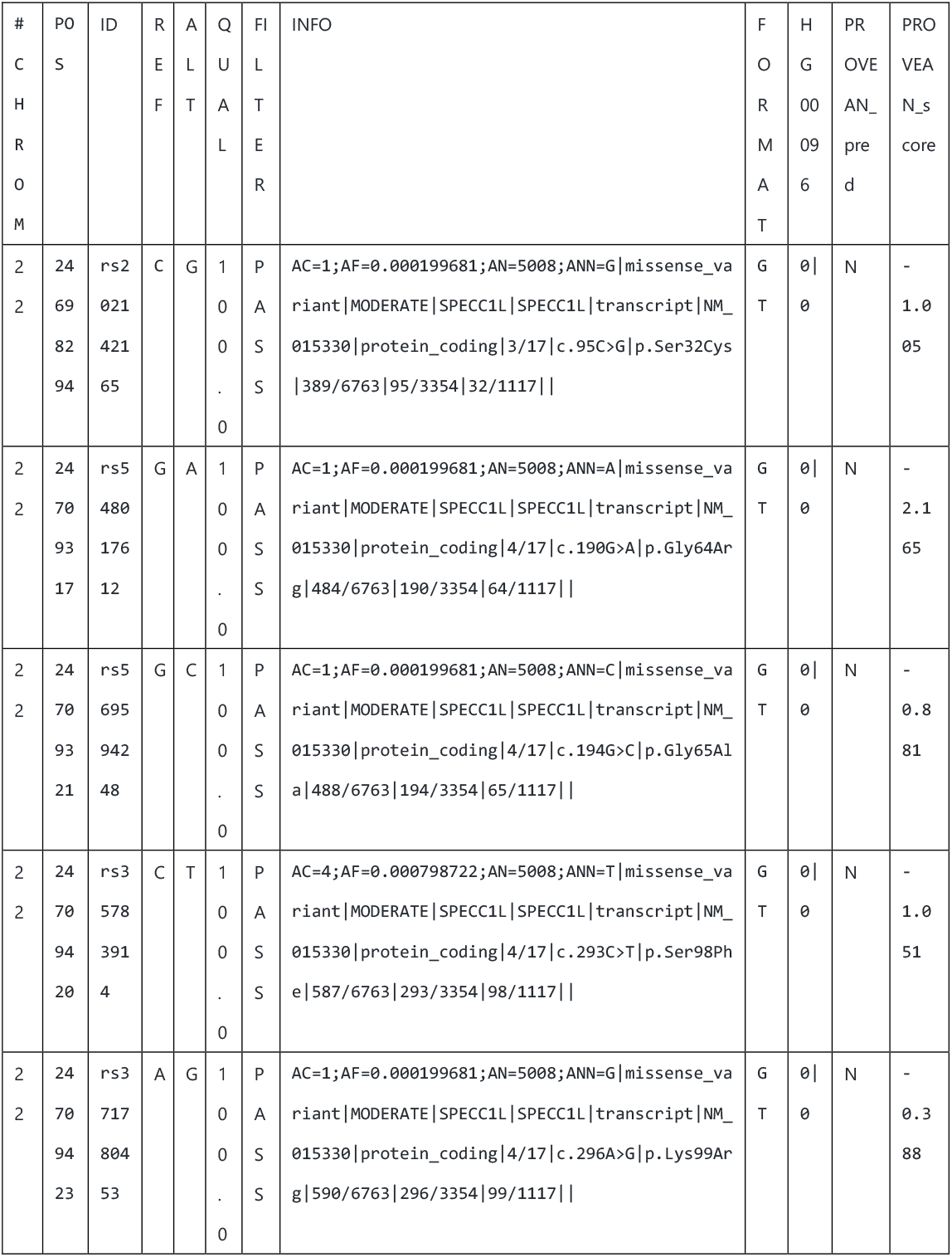
Output overview of ePat

Lists of SNVs/INDELs with rough PROVEAN annotations for each family ware created as before [6]. Of the number of SNVs/INDELs in this list that were not annotated by PROVEAN, we measured how many SNVs/INDELs were annotated with ePat. We created 1216 files and detected 491.5 SNVs/INDELs per family, of which 189.54 SNVs/INDELs per family were not annotated with conventional PROVEAN. Of the 189.54 SNVs/INDELs that were not annotated, we were able to annotate 151.63 SNVs/INDELs per family using ePat (Table 3).

**Table 3.** Statistical information of 1216 families used in the analysis

|  | average | rate[%] |
| --- | --- | --- |
| The Number of SNVs/INDELs | 491.5 | 100 |
| The Number of SNVs/INDELs annotated with PROVEAN | 301.95 | 61 |
| The Number of SNVs/INDELs not annotated with PROVEAN | 189.54 | 39 |
| The Number of SNVs/INDELs newly annotated with ePat | 151.63 | 31 |
| The Number of SNVs/INDELs annotated with ePat | 453.58 | 92 |

In the past, PROVEAN was only able to annotate 61% of the total SNVs/INDELs, and the rest of the data was not usable. Therefore, we developed ePat, which can calculate the toxicity of indel mutations with frameshift and mutations near splice junctions. As a result, we were able to annotate 92% of the total SNVs/INDELs by using ePat. In conclusion, ePat can help extract variants that affect biological functions with not only point mutations, and indel mutations that does not cause frameshift, but also frameshift, stop gain, and splice variants.

## 4. Software and code availability

The source code of ePat and the documentation is available from https://github.com/itotaku1225/ePat.

The singularity image and test data is available from https://zenodo.org/record/5482094#.YZSJtWDP2Uk.

